# Automated cryo-EM sample preparation by pin-printing and jet vitrification

**DOI:** 10.1101/651208

**Authors:** Raimond B.G. Ravelli, Frank J.T. Nijpels, Rene J.M. Henderikx, Giulia Weissenberger, Sanne Thewessem, Abril Gijsbers, Bart W.A.M.M. Beulen, Carmen López-Iglesias, Peter J. Peters

**Affiliations:** The Maastricht Multimodal Molecular Imaging Institute (M4I), Division of Nanoscopy, Maastricht University, Maastricht, the Netherlands; CryoSol World, Maastricht, the Netherlands; Instrument Development, Engineering and Evaluation (IDEE), Maastricht University, Maastricht, the Netherlands

**Keywords:** sample preparation, vitrification, VitroJet, cryo-electron microscopy, single particle analysis

## Abstract

The increasing demand for cryo-electron microscopy (cryo-EM) reveals drawbacks in current sample preparation protocols, such as sample waste and lack of reproducibility. Here, we present several technical developments that provide controlled and efficient sample preparation for cryo-EM studies. Pin printing substantially reduces sample waste by depositing only a sub-nanoliter volume of sample on the carrier surface. Sample evaporation is mitigated by dewpoint control feedback loops. The deposited sample is vitrified by jets of cryogen followed by submersion into a cryogen bath. Because the cryogen jets cool the sample from the center, premounted autogrids can be used and loaded directly into automated cryo-EMs. We integrated these steps into a single device, named VitroJet. The device’s performance was validated by resolving 4 standard proteins (apoferritin, GroEL, worm hemoglobin, beta-galactosidase) to ~3 Å resolution using a 200-kV electron microscope. The VitroJet offers a promising solution for improved automated sample preparation in cryo-EM studies.

## INTRODUCTION

Within just a few years, single-particle cryo-electron microscopy (cryo-EM) has become a powerful mainstream technique to resolve high-resolution 3D structures of macromolecules. Technological breakthroughs in microscopes, detectors, and processing have contributed to the resolution revolution^1^. However, numerous advancements are yet to be made^2, 3^. Reproducible sample preparation has emerged as one of the main bottlenecks^4^. The sample preparation technique for cryo-EM, pioneered by Dubochet and others^5–8^, allows particles to be preserved in a thin vitreous layer in their native state. The current leading commercial sample preparation devices still rely on the work of these pioneers. Sample preparation begins with an EM grid that is plasma-treated in a separate instrument to increase hydrophilicity. As this effect is only temporary, the wetting properties depend on the time between glow discharge and sample deposition, temperature, humidity and cleanliness of the environment^9–11^. Next, a few microliters of sample are manually pipetted onto the grid, and >99.99% of it is blotted away by filter paper, after which an aqueous film is allowed to thin by wicking and evaporation. During this dynamic process, air–water interfaces will form^5, 12, 13^, which can be detrimental to the structure of interest as proteins tend to absorb to such interfaces and (partly) denature^12^. The resulting film of sample on a perforated carrier^14–16^ has a generally concave shape^12, 17^ due to drying and draining, with the center being the thinnest. The grid is held by tweezers and plunged into a bath of cryogen to vitrify the sample, so that it can be observed in the vacuum of the microscope. This vitrification should be fast enough to prevent the formation of ice crystals. Grids are stored under cryogenic conditions until analysis. For microscope loading and unloading, the fragile grid needs to be assembled into either a sturdy cartridge or the tip of a holder: both processes are cumbersome and often harmful to the fragile grid.

Many steps of the above protocol require skilled operators, careful handling, clean tools, clean cryogens, and controlled environments. Some steps are intrinsically difficult to control accurately, such as positioning of the filter papers, blotting force, humidity and flatness of the filter papers, as well as the position and shape of the tweezers. As a result, reproducibility is lacking. The required training and skills to obtain reliable grid quality is a significant entry barrier for new scientists. The increasing demand and evolution of cryo-EM call for improved, scalable methods. We have developed a more controlled method with minimal operator intervention by employing some steps new to the field of cryo-EM and automating the entire workflow. Our method consists of (i) an integrated glow-discharge module to control and minimize the time between plasma cleaning and sample deposition; (ii) pin-printing for sample application, which requires only sub-nanoliter sample volumes and eliminates sample blotting; and (iii) jet vitrification, which allows for the handling of autogrids. We integrated these features into a single setup, termed the VitroJet, and used it to prepare four standard proteins to obtain high-resolution single particle reconstructions.

## RESULTS

### Sample carriers

All results below were obtained with pre-mounted autogrids^18, 19^. Autogrids were initially developed for increased robustness in order to allow automated handling of grids in cryo-transmission electron microscopes (TEMs), such as the Titan Krios, Glacios and Arctica (Thermo Fisher Scientific). Traditional vitrification devices are not able to vitrify pre-mounted (assembled at room temperature) autogrids. Therefore, autogrids are normally manually assembled under cryogenic conditions by clamping an EM grid into a sturdy cartridge by means of a flexible C-clip spring (so called post-mounting). The jet vitrification procedure described below overcomes this limitation.

### Glow discharge module

Traditional vitrification devices use external glow discharge modules. We characterized this procedure by determining contact angles for typical glow discharge settings used in our lab. An ELMO Glow Discharge System (Cordouan Technologies) was operated at 7 mA, 0.35 mbar, and 30 s glow discharge time. Contact angles (measured using a Krüss drop shape analyser, model DSA25) of 25 degrees and less were obtained for minimal transfer times (<1 min) between the glow discharge system and the drop shape analyser.

Within the VitroJet, the glow discharge module is an integral part of the device. Each grid is glow discharged separately, and transferred within 10 s from the glow discharge unit into the process chamber. We evaluated different glow discharge settings (0.3–20 mA; 0.05–0.2 mbar), using different grids (Quantifoil R2/2 Cu300, UltraAuFoil R2/2 Au300), different treatment times (5–40 sec), and different durations between the glow discharge treatment and the contact angle measurement (1–17 min). Excellent contact angles from grids glow-discharged in the VitroJet could be obtained. In general, longer glow discharge treatment gave lower contact angles. Not surprisingly, longer delays (17 min) between glow discharge and contact angle measurements gave larger contact angles (35 degrees). The glow-discharged grids did not show any visible damage (data not shown). The data below were obtained using 30 s glow discharge at 0.5 mA and 0.1 mbar (yielding contact angles <25 degrees).

### Process chamber

In order to minimize sample evaporation, current vitrification devices such as the Vitrobot, Leica EM GP, and the Gatan Cryoplunge apply the sample inside a humidified chamber. Passmore^9^ recommended to keep the relative humidity (RH) surrounding the specimen support at 100% to prevent changes in the solute concentration prior to freezing. However, it is difficult to achieve a reliable 100% RH. Once the air within the chamber is fully saturated with water vapour, condensation will occur. Water droplets on the humidity sensor and/or the grid compromise the reproducibility of the experiment: humidity sensors only work reliably under non-condensing conditions.

We first studied the theoretical and reported evaporation rates of thin layers of water on grids. Different evaporation velocities of thin (suitable for TEM) water layers have been reported, depending on temperature and RH: 5 nm/s at 4 °C and 90% RH^9^ and 40–50 nm/s at 20 °C at 40% RH^13^. We modelled the evaporation of a thin layer of water on a grid (supplementary materials). Parameters within this model include partial water pressure in air, saturated vapour pressure, velocity of the air flow over the sample, thermal conductivity of the sample carrier, and shape of the deposited water layer. For a given temperature and RH, the dewpoint temperature of the grid can be calculated (Figure 1a). At <100% RH, the dewpoint temperature of the grid will be lower than the temperature in the chamber. Our model predicts that the layer thickness decreases by 70 nm/s for every °C of dewpoint error, for a thin line of pure water deposited onto the grid at 93% RH in the chamber (Figure 1b; supplementary materials).

**FIGURE 1.**
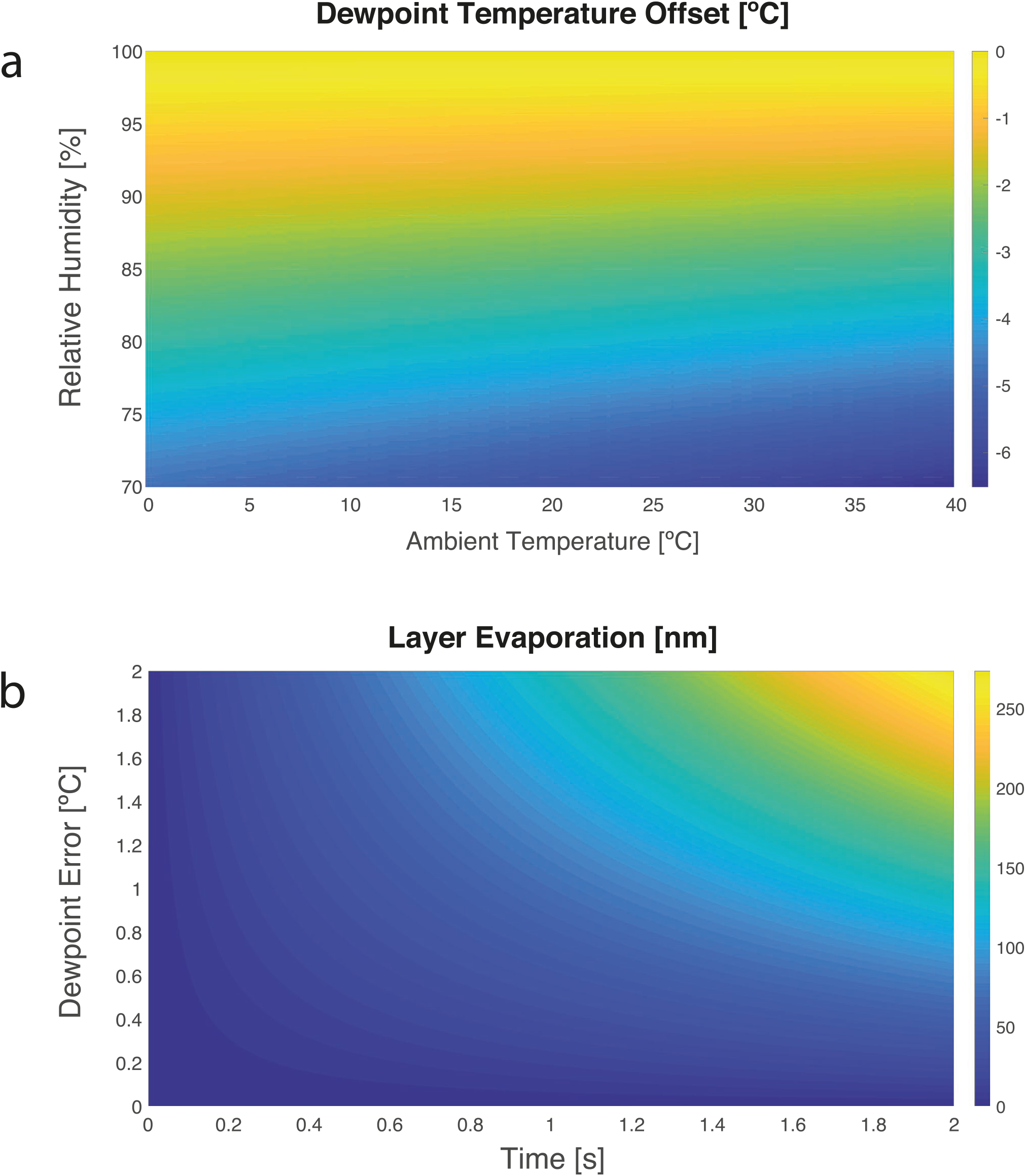
Dewpoint temperature calculations and predictions. (a) Dewpoint temperature offset (color scale, in degrees) as a function of the relative humidity and ambient temperature. *E.g*. when working at 20 °C and 94% relative humidity, the dewpoint temperature would be 1°C (orange) below the ambient temperature. (b) Predicted decrease in thickness of a thin sample layer due to evaporation (color scale, in nm/s) with respect to the time and the dewpoint error, which is the difference between sample temperature and its dewpoint. *E.g*. for a temperature of 1 °C above dewpoint, 70 nm of the layer is estimated to evaporate within one second.

In order to prevent sample evaporation, the sample itself thus needs to be maintained at dewpoint. We built a fast feedback loop to maintain the grid at dewpoint temperature while monitoring the RH and temperature of the process chamber. Within the VitroJet, this chamber is maintained at 93% RH. The grid is peltier-cooled through the autogrid cartridge. The optimal grid temperature was offset from its calculated dewpoint to account for heat exchange between the grid and its environment, resulting in an elevated temperature of the center of the grid compared to the sensor positioned just below the grid. Even if the edge of the grid would be held perfectly at the dewpoint temperature, the estimated temperature of the center can be a few tenths of a degree higher (supplementary materials). Our theoretical model predicts that, if such temperature difference would not be accounted for, the layer thickness could still decrease by 14 nm/s. An incorporated camera enables visual inspection of evaporation and/or condensation on the grid and can be used to finetune the feedback loop.

### Pin printing

To improve sample deposition on EM grids, several developments have been presented, including droplet-based methods^20–24^, *e.g*. on nanowire grids^25^, and using capillaries followed by sample thinning by controlled water evaporation^26^. We sought a way to obtain sample thickness layers suitable for cryo-EM without the need for blotting, nanowire grids, and/or extra water evaporation steps. In our pin-printing method, a capillary bridge is formed between a metal pin and the EM grid, holding a sub-nanoliter volume of sample. As the bridge is moved across the surface of the carrier (be it modified or off the shelf), it leaves behind a sample layer of uniform thickness suitable for cryo-EM SPA studies. A stock sample volume of 0.5 μl is introduced into the process chamber by a positive displacement pipette. A metal pin is dipped into the sample to collect a sub-nanoliter volume. The pin is cooled down to dewpoint to prevent evaporation of the tiny droplet at its tip. The pin is moved to a predefined distance (typically 10 µm) from the carrier surface such that the sample forms a capillary bridge between the pin and grid. As EM grids do not have a perfectly flat surface, the absolute reference positions of the pin and autogrid are calibrated separately in the VitroJet using the camera. Once the capillary bridge is formed, the pin is moved along the surface of the carrier (typical speed 0.3 mm/s; Supplementary movie 1). Capillary forces ensure that the liquid bridge follows the pin. If this parallel movement is sufficiently fast, a thin film will be deposited due to the viscous shearing on the liquid. The film thickness *h* is determined by the relative speed of the pin *u* moving over the carrier surface, the stand-off distance *δ* between pin and carrier, the viscosity *μ*, surface tension σ, and surface properties of the pin and the carrier (Figure 2a). By altering these parameters, variations in fluid properties can be counteracted and the deposited layer thickness can be tuned.

**FIGURE 2.**
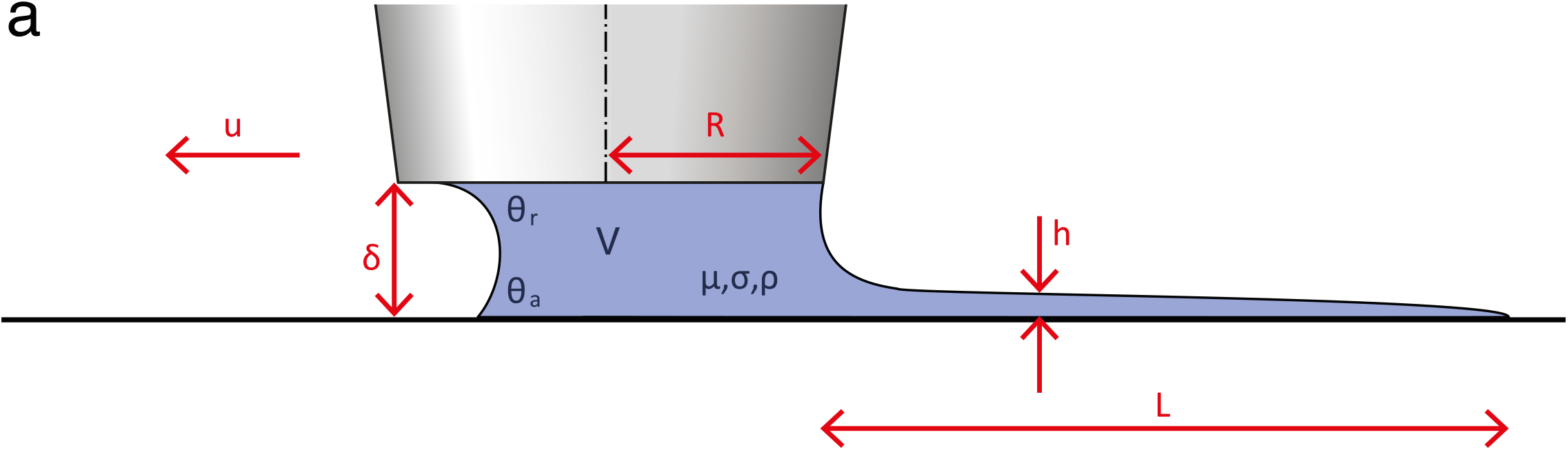
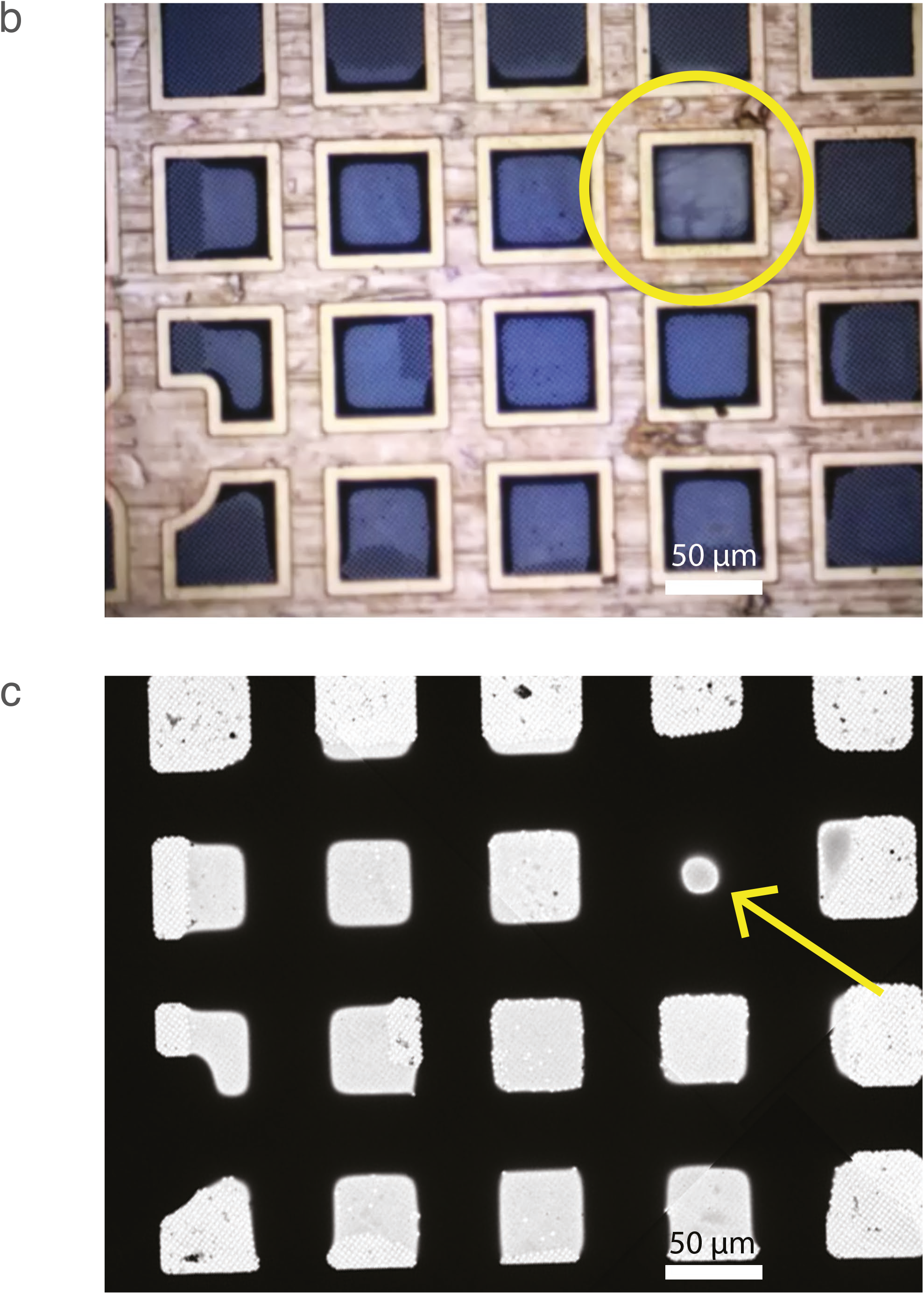
Pin printing parameters and resulting sample deposition. (a) Schematic overview of relevant parameters for pin printing. The deposited layer in height (*h*) and length (*L*) is a result of the sample properties (viscosity *μ*, surface tension *σ*, density *ρ*), contact angles (advancing at the grid *θ*_a_, receding at the pin *θ*_r_), and writing parameters (standoff distance *δ*, pin radius *R*, pin velocity *u*). (b) Photograph of the grid by the integrated camera shows the sample applied by pin printing. The pin is indicated by a yellow circle. (c) Low magnification cryo-EM atlas of the deposited and vitrified sample. One thicker square (arrow) is the result of retraction of the pin from the grid. Scale bars represent 50 μm.

While printing could be performed in any pattern, we found that printing in a partly overlapping spiral pattern resulted in a more uniform thickness compared to, for example, in one straight line across the carrier. Finally, the pin retracts from the surface and the remaining volume in the liquid bridge breaks up, leaving a thicker droplet at the end of the deposited line (Figure 2b, c). Because only a sub-nanoliter volume is deposited per printing, numerous grids can be printed from the same stock volume of 0.5 μl.

The initially deposited film layer could differ in thickness from the final one that is vitrified. Possible reasons for this include evaporation, condensation, and liquid flow. The first two are minimized by tight control of the dewpoint feedback loop described above. Liquid flow in the deposited thin film takes place by wicking. As the grid itself is wetting, the crevices between perforated foil and grid bars will absorb sample through capillarity. Sample absorption into these crevices is limited by the flow within the thin layer. Typical time scales are given by the redistribution time scale *T*_*r*_ for a surface-tension-driven thin-film flow^27^: *T*_r_ = *μd*^4^/*σh*^3^, where *μ* ≈ 10^−3^ *Pa s* is the dynamic viscosity of the liquid, *d* = 50 *μm* is the distance between the center of a grid square and the grid bar, *σ* ≈ 0.07 *N*/*m* is the surface tension of the liquid, and ℎ is the thickness of the liquid layer (e.g. 50nm). In such case, the redistribution of sample from the center of the square would take >10 minutes. For mesh sizes that have larger grid squares or thinner deposited layers, this time scale will become even larger. Only the area that is very close to the grid bar will be affected by wicking-induced liquid flow, since vitrification halts the layer thinning directly after sample deposition. The thin liquid layer within each grid square remains intact (figure 2c), illustrating that evaporation and liquid flow within these thin layers are minimal.

### Vitrification module

After sample deposition, traditional vitrification devices rapidly plunge the bare grid into a bath of cryogenic liquid to vitrify the sample. Plunge vitrification starts cooling down from the bottom of the grid upwards, and boiling of the coolant can form a gaseous insulating layer at the surface of the grid. This process can compromise the cooling time of the sample, which is estimated to be 10^−4^ s in the most favourable case of thin-layer vitrification^7^. These current devices are not able to vitrify pre-mounted autogrids due to the extra thermal mass of the cartridge. The sturdy rim of autogrids would hit the cryogenic liquid first, whereas the area of interest (the sample on the middle of the grid) is cooled later at speeds too slow to prevent ice crystal nucleation.

Inspired by cryofixation of thick tissues for room-temperature ultrastructural studies^28^, we devised an alternative way to vitrify samples in pre-mounted autogrids: jet vitrification. Two streams of cryogenic liquid hit the sample and its carrier in the center (Supplementary movie 2), cooling it down to temperatures below 130 K in <1 ms. As jet vitrification cools the autogrid from the center outwards, the rim of the cartridge is cooled down last. The jets continues to spray for 50 ms to precool the cartridge; hereafter, the autogrid is submersed into a bath of liquid ethane to fully cool down the cartridge rim and gripper. The grid is subsequently slowly moved out of the ethane bath to allow excess ethane to flow off and prevent ethane solidification. Following vitrification, the gripper transfers the autogrid to a spring-loaded storage container in liquid nitrogen to connect directly to the microscope-loading workflow.

We conducted experiments with different cooling media at different temperatures, including liquid nitrogen (77 K), ethane/propane (37%/63% (v/v) at 79 K and 93 K), and ethane (99 K). Of these different media, liquid nitrogen and liquid ethane/propane cooled the grid to the lowest final temperature, but the highest cooling rates were obtained with liquid ethane (Supplementary Figure 1). These cooling rates were measured with 25 μm constantan wires (Bare thermocouple wire, Omega, Norwalk) woven into a copper grid mesh to form a thermocouple: such wires are expected to cool much slower than the very thin sample. The center of the grid showed significantly higher cooling rates with jetting compared to plunging, both for autogrids as well as regular grids (Supplementary figure 1).

### Integration

The above steps (glow discharge, process chamber, pin printing, and jet vitrification) were implemented into the VitroJet (Figure 3a). A supply cassette provides up to 12 pre-clipped autogrids. A gripper picks up each autogrid individually and transports it sequentialy through each of the different steps for sample preparation (Figure 3b; Supplementary movie 3). The gripper is dried by nitrogen gas within the glow discharge unit before picking up the next carrier.

**FIGURE 3.**
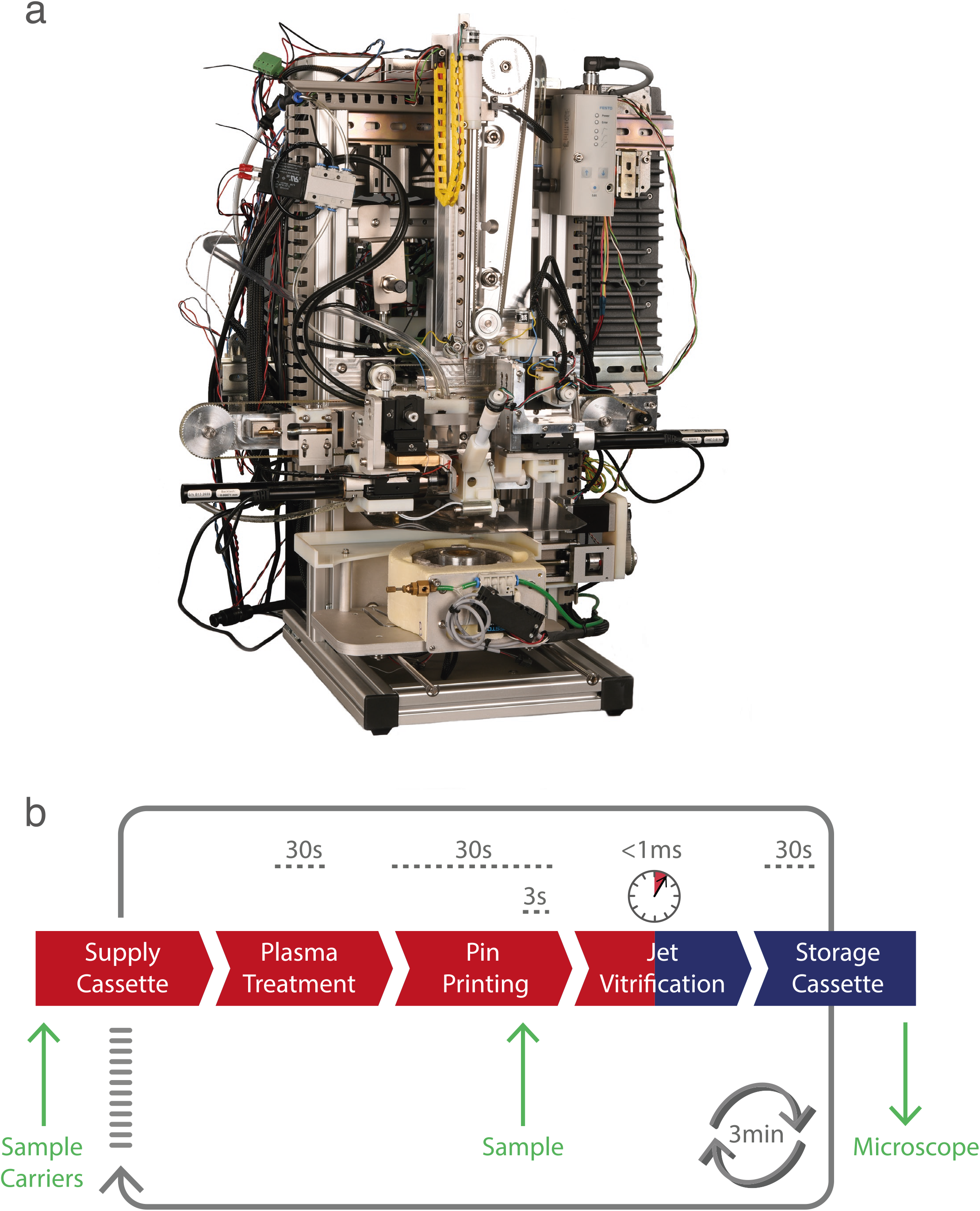
The VitroJet workflow. (a) The device and (b) the workflow. Sample carriers (up to 12) are introduced in a supply cassette and sample (up to 0.5 µl) in a pipette tip (yellow arrows). The VitroJet automatically processes the cassette by sequentially passing sample carriers through the plasma treatment, sample deposition by pin printing, jet vitrification and finally storing the grid in a cryogenic cassette ready to enter the electron microscope (green arrow). The supply cassette is at room temperature (red), the storage cassette at cryogenic temperature (bleu). An animation showing all subsequent steps is shown in Supplementary movie 1.

The cycle time of the workflow is ~3 min. Most of this time is taken up by the glow discharge module: evacuation of this chamber takes approximately 20 s (twice per cycle), while the glow discharge itself takes 30 s. For pin printing, alignment of the sample carrier and pin within the process chamber requires < 30 s, and the sample deposition a few seconds. Transfer of the grid between the process chamber and the vitrification chamber is completed within 80 ms. Vitrification of the sample lasts < 1 ms, whereas another 300 ms is used to deeply cool down the entire autogrid assembly. Grid removal from the ethane bath and transfer requires < 30 s.

Before each use, the VitroJet undergoes several automatic preparatory tasks, which are completed in ~15 min and includes cooling down the vitrification unit, filling the liquid ethane reservoir, and equilibrating the process module at the right humidity. The temperature-controlled liquid ethane reservoir is cooled by a liquid nitrogen bath.

### Structure determination by single particle analysis (SPA)

To validate the VitroJet, we prepared samples of several standard proteins (apoferritin, GroEL, worm hemoglobin, beta-galactosidase) and performed high-resolution SPA (see methods). Each sample was pin-printed on *~*16 squares of 300-mesh grids with perforated foils (R1.2/R1.3) and jet-vitrified using liquid ethane. Atlas overviews collected within the cryo-EM show excellent correlation with the visualization of the deposition within the VitroJet just prior to vitrification (Figure 2bc). Cryo-EM data were collected on a 200-kV FEI Arctica microscope. From the squares that were pin-printed, most holes could be selected for data collection. Holes close to the grid bar were skipped to avoid thicker ice due to the wicking of the grid bars. Tomographic inspection of different holes within the bulk of one square showed consistent ice thickness of ~40 nm. Micrographs were recorded between 0.5 and 2.0 μm underfocus, showing good contrast (Figure 4).

**FIGURE 4.**
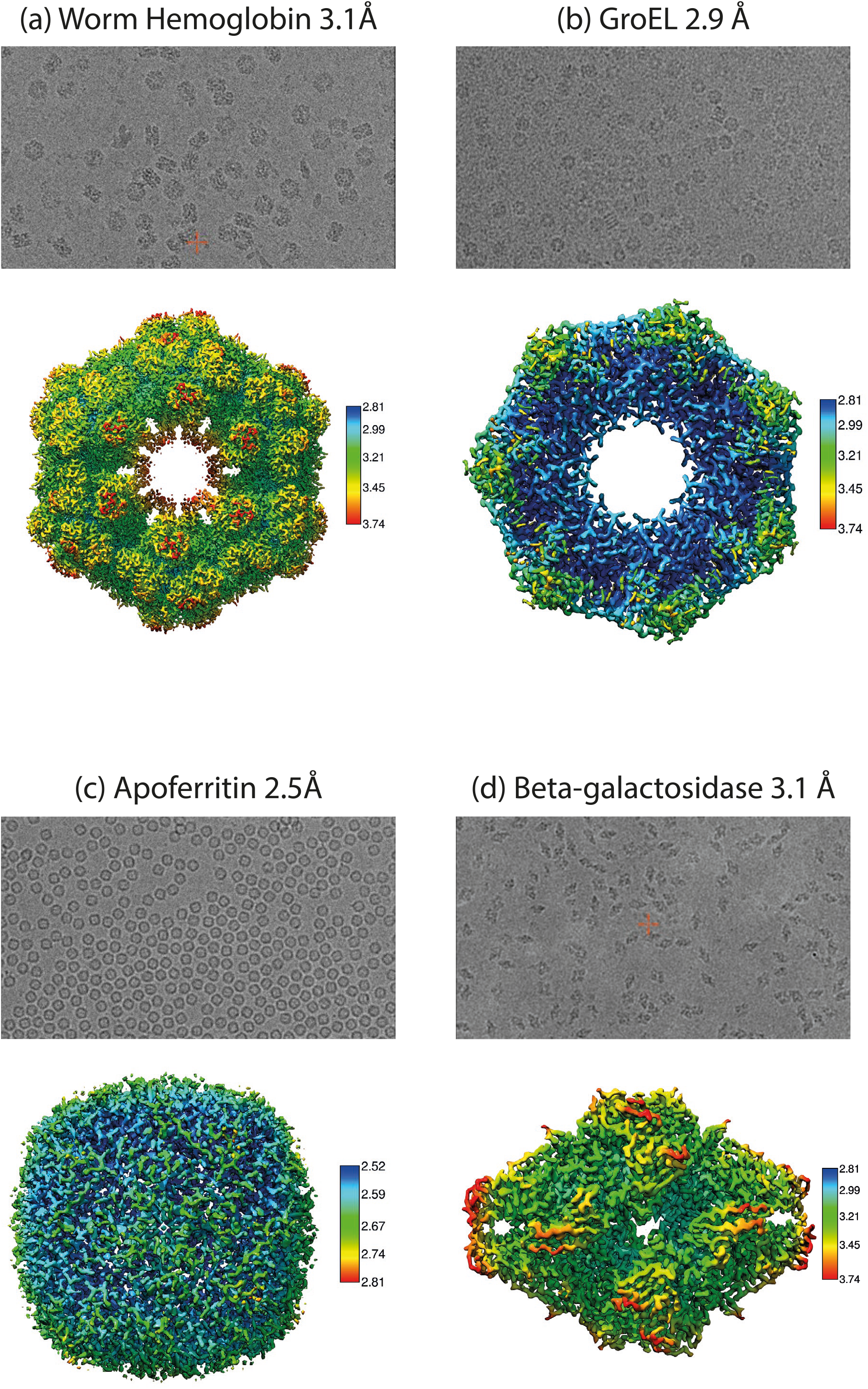
Sample preparation of four standard proteins by the VitroJet, and validated by cryo-EM analysis. Micrographs (top) and reconstructions (bottom) of worm hemoglobin, GroEL, apoferritin, beta-galactosidase. The reconstructions are coloured according to local resolution as calculated using Relion.

The data, processed with Relion^29^, yielded good 2D classes. After iterations both with particle picking and extraction, (local) contrast transfer function estimation, 2D and 3D classifications, 3D reconstructions of ~3 Å resolution were obtained for each of these four proteins (Figure 4). Apoferritin gave a 3D reconstruction of 2.49 Å using 47209 particles (O symmetry). For GroEL, we obtained a 3D reconstruction of 2.94 Å using 9809 particles (D7 symmetry). 3D reconstructions of worm hemoglobin and beta-galactosidase had respective resolutions of 3.11 Å (using 10488 particles, D6 symmetry) and 3.11 Å (using 15252 particles, C7 symmetry). Models refined against the four reconstructions were in accordance with earlier published models demonstrating the success of the VitroJet in preparing suitable samples for cryo-EM studies.

## Discussion

Traditional cryo-EM sample preparation methods require multiple manual steps and depend on ill controlled parameters such as blotting force, grid positioning, and time between glow discharge and sample application. Here we present a workflow that minimises operator dependency and provides control over all relevant parameters. Prior to starting the sample preparation cycle, parameters regarding glow discharge, dewpoint, pin printing and jetting can be set. After initiation, the process is executed in an automated fashion. In an era where automation has made so much impact on microscope alignment, data collection, processing and model building, automated control over sample preparation is a mandatory next step.

Samples applied to pre-mounted autogrids are difficult to blot and even more difficult to vitrify using the existing leading commercial devices. We overcame both problems by using pin printing, which does not require blotting, and jet vitrification, which yields superior cooling rates starting from the center of the grid where the sample is located. In addition to circumventing the problems associated with blotting, pin printing requires minute volumes of sample, which enables the study of macromolecules for which only micrograms can be obtained.

Proteins tend to absorb to the air–water interface, referred to as “the deadly touch”^12^. It seems intuitive that reducing the time the sample is exposed to such an interface would help to prevent protein denaturation^17^. However, using the Stokes Einstein equation, it has been calculated that even for a minimal residence time of ~1 ms, particle-surface interactions will occur dozens of times before the water is frozen^30, 31^. The VitroJet has minimal transit time between sample application and vitrification (80 ms); however, such times are still long compared to the minimal residence time. A promising complementary approach would be to control the surfaces present in the specimen support^30, 32^, as demonstrated for yeast fatty acid synthase applied on a substrate of hydrophilised graphene^12^. The pin printing procedure presented here is compatible with a multitude of (modified) grid supports.

Jet vitrification was originally demonstrated 40 years ago, on thick tissue spanned over a holder^28^. The method was used as a prelude to freeze substitution, resin embedding and sectioning, for ultrastructural studies performed at room temperature. Sample evaporation was not a concern for the bulky tissues used. Here we adapted jet vitrification to obtain thin layers of macromolecular samples. One might expect that sample evaporation problems would be insurmountable and that the jets would blow away the thin liquid layers of macromolecules prior to vitrification, resulting in empty holes. However, we have demonstrated the opposite. We believe that the sample already vitrifies before the liquid cryogen hits the sample^33^.

The setup of the VitroJet is modular, making it easier to incorporate future developments to further advance the cryo-EM field. The pin printing applies sample to a part of the available area of the grid future generations of the VitroJet could be equipped with multiple pins moving simultaneously over different parts of the grids. Such schemes would enable higher-throughput screening as well as time-resolved studies^34, 35^, e.g. combined with laser excitation. The process chamber provides the ability to control condensation as well as evaporation for an extended duration of time on a specific layer thickness, which could also offer benefits for soft condensed matter studies. While the VitroJet described here was developed for SPA, we aim to develop a branch of the VitroJet dedicated to the preparation and vitrification of cellular samples. Vitrification of cells is inherently more difficult than that of purified macromolecular samples. For example, it was stated that the center of HeLa cells clearly undergoes incomplete vitrification^36^. Preliminary results indicate that jet vitrification will help to reduce this problem, which would be a true asset for *in situ* structural biology. The VitroJet offers much-needed innovations in sample preparation, which will accelerate and perhaps even revolutionize future cryo-EM studies.

## Methods

### Protein purification

Human apoferritin overexpressed in *E. coli* was kindly provided via Evgeniya Pechnikova (Thermo-Fisher) by Dr. Fei Sun (Institute of Biophysics, Chinese Academy of Sciences). Chaperonin-60 from *E. coli* (GroEL) was ordered from Sigma (C7688), dissolved in a buffer comprised 50 mM Tris (pH 8.0), 100 mM KCl, 10 mM MgCl_2_, 2 mM DTT and 80 mM trehalose, and used without further purification at a concentration of 10 mg/ml. ~400 μl Blood from earthworm *Lumbricus terrestris* (WHBG) was extracted from the seventh segment of the body^37^ and injected into a Superose® 6 Increase 10/300 GL size-exclusion chromatography column (SEC) with an elution buffer of 20 mM Tris (pH 8.0), 150 mM NaCl, 10 mM CaCl_2_. After purification, WHBG was concentrated to 6 mg/ml. Beta-galactosidase (β-gal) from *E. coli*, ordered from Sigma (G5635), was further purify by SEC with a Superdex® 200 Increase 10/300 GL column and eluted with 25 mM Tris (pH 8.0), 50 mM NaCl, 2 mM MgCl_2_ and 1 mM TCEP. The ligand phenylethyl β-D-thiogalactopyranoside (PETG) was purchased from Sigma-Aldrich (catalog #P1692) and prepared as described^38^. We used a final β-gal protein concentration of 4 mg/ml with 7.5 mM PETG.

### Sample preparation

Quantifoil R1.2/1.3 Au300 and UltraAufoil R1.2/1.3 Au300 grids (Quantifoil Micro Tools, Jena, Germany) were used as sample carriers. Grids were pre-clipped before entering the VitroJet. Sample deposition occurred in a climate chamber at room temperature with a humidity of 93%, where pin and grid were cooled towards dewpoint temperature. A sub-nanoliter volume of sample was pin-printed onto the grids with a writing speed of 0.3 mm/s. Samples were vitrified within 80 ms after sample deposition by two pressurized jets of liquid ethane for 50 ms at 99 K and 1 bar.

### Single particle cryo-EM

Micrographs were collected on a 200-kV Thermo Fisher Tecnai Arctica microscope equipped with a Falcon3 detector. For each standard protein, micrographs were collected with a calibrated pixel size of 0.935 Å. For apoferritin and GroEL, total integrated electron flux of ~40 e^−^/Å^2^ in counting mode at a defocus range of 0.7–1.5-μm underfocus was used. For WHBG and β-gal, a total integrated electron flux of ~43 e^−^/Å^2^ in counting mode, at 0.6–1.4-μm underfocus for WHBG and 0.5–1.3-μm underfocus for β-gal, was used. We recorded 383 movies over 48 s for apoferritin, 1284 movies over 69 s for GroEL, 1115 movies over 77 s for WHBG, and 352 movies over 78 s for β-gal.

### Image processing

The images were processed in Relion^39^, where the frames of the movies were aligned and averaged using a Bayesian approach as described^40^. The contrast transfer function (CTF) parameters were calculated with GctF^41^. Afterwards, a subset of micrographs was used to pick ±500 particles manually for initial 2D classification. These 2D classes were used for an iterative, automated particle-picking procedure where both the references and the autopicking parameters were improved using a subset of the micrographs. The complete data set was autopicked, particles were extracted and subjected to an iterative 2D classification scheme to reject bad particles. After 2D classification, 3D auto-refine was performed using starting models based from the PDB (4W1I, 2YNJ, 1×9F, 6CVM). The resulting reconstructions were low pass filtered and projected for another iteration of picking, 2D classification, and 3D auto-refine. Local CTF refinement, and local symmetry in the case of WHGB, resulted in the final maps. Given resolutions were estimated based on established, Fourier Shell Correlation^42^ standards^43^, as directed by the program Relion.

### Model refinement

The PDB starting models mentioned above (4W1I, 2YNJ, 1×9F, 6CVM) were superimposed on the sharpened cryo-EM maps. Models were refined iteratively through rounds of manually adjustment in Coot^44^, real space refinement in Phenix^45^ and structure validation using MolProbity^46^. Data have been deposited in the EMDB (deposition codes X1 X2 X3 X4) and PDB (deposition codes Y1 Y2 Y3 Y4).

## Author contributions

PJP, CLI and RBGR designed and directed the project; FJTN designed and constructed the machine with input from all authors and BWAMMB as project leader; ST did preliminary tests; RJMH and GW performed the experiments; AG prepared the samples; RBGR, RJMH and GW analysed the data; RBGR and RJMH wrote the manuscript with input from all authors.

## Acknowledgements

We thank Dr. Fei Sun (Institute of Biophysics, Chinese Academy of Sciences) for providing apoferritin sample, Pascal Huysmans and Paul Kwant (IDEE, Maastricht) for engineering input, Peter Frederik for helpful discussions and Hang Nguyen for critical reading of the manuscript. Hans Duimel and Hirotoshi Furusho provided technical support from the UM Microscopy Core Lab, Giancarlo Tria helped with initial experiments, Paul van Schayck with the IT infrastructure, and Roger Jeurissen with the theoretical framework. This research received funding from the Netherlands Organisation for Scientific Research (NWO) in the framework of the Fund New Chemical Innovations, numbers 731.014.109 and 731.016.407, as well as from the Province of Limburg, the Netherlands.

## Statement about competing financial interests

The University of Maastricht filed patents with some of the authors as inventors regarding sample preparation for cryoEM as outlined in this manuscript. PJP is shareholder and CSO of the startup CryoSol-World that holds the licensees of these submitted patents.

## Supplementary figures

Supplementary Figure 1. Cooling rates of sample carriers by the different vitrification methods. Measurement of average rate (K/s) at which the thermocouple is cooled down from 273 K to 123 K. The thermocouple consists of constantan wire woven in a copper grid, which adds extra thermal mass to the grid and therefore influences the measurement. As the biological samples on the thin films are located away from the grid bars, these measurement is used as surrogate marker for the different cooling methods. Ethane jet vitrification results in significant higher cooling rates compared to ethane plunge vitrification. Similar cooling rates were found regardless of the jet vitrification pressure (1–3 bar). For all methods, liquid ethane (red diamonds) showed the highest cooling rates compared to their mixtures with propane (green and blue symbols) or to liquid nitrogen alone (orange).

Supplementary movie 1. Animation showing the working principles of the VitroJet.

Supplementary movie 2. Video recorded by the grid inspection camera showing the sample deposition in real time. The presence of a suitable thin layer as well as the occurrence of sample evaporation or condensation can be assessed.

Supplementary movie 3. Slow motion video recorded by a high speed camera showing vitrification of an autogrid with two jets of liquid ethane. The jets last for 50 ms.

