## Supplementary Materials for "Automated cryo-EM sample preparation by pin-printing and jet vitrification"

**Evaporation**

Evaporation of sample will lead to a decrease in layer thickness, concentration of buffer components and changes in osmotic pressure. In cryo-EM sample preparation, evaporation will play a significant role due to large surface to volume ratio's before vitrification. Inside the VitroJet the sample is cooled down to its dewpoint temperature to minimize evaporation. This supplementary material provides an order of magnitude estimation of the evaporation rate depending on the error from its dewpoint temperature. As there exists a temperature variation over the grid itself, evaporation will be dependent on the location on the grid.

The evaporation rate will be estimated as a function of the offset in grid temperature from its dewpoint. The dewpoint temperature can be calculated according to the following equation:

$$T_{dp} = \frac{c\gamma(T, RH)}{b - \gamma(T, RH)}$$

where $b = 18.678$ and $c = 257.14\ °C$ are constants, whereas $\gamma$ is a function of the temperature $T$ (in degrees Celsius) and the relative humidity *RH.*

$$\gamma = \ln\left(\frac{RH}{100}\right) + \frac{bT}{c + T}$$

When working at 20 °C and 93% humidity, the dewpoint is 1.2 $°C$ lower than the ambient temperature in the chamber.

The temperature of the grid is mainly dependent on two factors. Firstly, the grid is cooled down toward its dewpoint temperature by conduction. In order to do so, the copper autogrid ring is pressed against a temperature-controlled element. As the thermal mass of the autogrid ring is much larger than that of the grid, we treat the copper ring as a thermal reservoir. The EM-grid itself consists of a perforated carbon foil supported by a gold grid. Below we use grids with grid bars that are 10 μm thick, have a thermal conductivity of 314 W $m^{-1}$ $K^{-1}$, and a perforated carbon foil of 12 nm thickness with a thermal conductivity in the order of 1 W $m^{-1}K^{-1}$. This indicates that the thermal resistance of the carbon foil is large with respect to the thermal resistance of the gold grid. Therefore, heat conduction in such EM-grid occurs mainly through the grid bars.

Secondly, the air in the chamber will heat up the grid through convection. In this case, only natural convection will occur since deposition takes place in a closed chamber and the flow of the humidifier is turned off during deposition. The air flow in the chamber will not be forced, but depends on the temperature differences in the chamber. In order to determine the nature of natural convection we calculate the Rayleigh number. This dimensionless number gives the balance between the driving force (buoyancy) and viscous dissipation, and can be determined using the following formula

$$Ra = \frac{gL^3\Delta T}{\nu\alpha T}$$

The Rayleigh number depends on the gravitational acceleration $g = 9.81\ m\,s^{-2}$, the length scale $L = 3\ mm$ which is the grid diameter, the temperature difference $\Delta T = 1.2\ K$ between the sample and the air in the chamber, the kinematic viscosity $\nu = 1.4 \cdot 10^{-5}\ m^2s^{-1}$ of the air in the chamber, the thermal diffusivity of the air $\alpha = 1.1 \cdot 10^{-5}\ m^2s^{-1}$ in the chamber, and the absolute temperature $T = 293\ K$ of the air in Kelvin. The Prandtl number can be used to determine the thermal diffusivity ($Pr = \frac{\alpha}{\nu}$). When using these values, the Rayleigh number is Ra = 7, which shows that both thermal conduction and thermal convection are relevant. As the value is well below the critical Rayleigh number for the transition to turbulent flow ($Ra < 5 \cdot 10^4$), thermal convection is laminar[47].

The evaporation rate is depending on the diffusion rate of water through the air, and on the heat transfer. Diffusion of water can limit the evaporation by raising the relative humidity close to the deposited line. The heat transfer can limit the evaporation rate since the heat of evaporation must be replenished through the air and the grid. To determine which of the two is limiting, we calculate the Péclet number. This dimensionless number is determined by ratio of advective- over diffusive heat transfer:

$$Pe = \frac{u^2 w}{D}$$

The Péclet number is dependent on the length scale of the heat transfer which is the width of the deposited line $w = 200\ \mu m$, the mass diffusivity $D = 1.4\ 10^{-5}\ m^2 s^{-1}$, and the velocity $u$. The velocity of the air can be obtained from the thermally driven convection.

$$u = \frac{\alpha}{L} Ra^{\frac{1}{4}}$$

This allows us to determine the Péclet number, which is equal to 0.1. As this shows the ratio of advective over diffusive heat transfer, it shows that diffusion is dominant in this situation. Afterwards, the expression for the evaporation rate can be calculated through the Sherwood number, which indicates the dimensionless mass transfer. As the situation is diffusion limited, the Sherwood number is of the order of unity ($Sh \approx 1$). Using this Sherwood number, the mass flux per unit surface can be determined with the following equation

$$j = \frac{Sh \cdot D}{w}\frac{dc}{dT}$$

in which *j* is the molar mass flux of water by evaporation, *D* is the mass diffusivity and $\frac{\mathrm{d}c}{dT}$ is the concentration difference which is depending on the temperature. The value that should be used here is the concentration difference between the air directly in contact with the liquid, and the concentration of water at the temperature of the liquid. If the liquid temperature is exactly equal to the dewpoint, this concentration difference is zero. However, if the sample temperature is too low then the moisture will be depleted slightly, decreasing the dewpoint temperature. When the temperature rises again to the original dewpoint, liquid will evaporate because the dewpoint has decreased. If the temperature is too high, liquid will evaporate even if the dewpoint had not changed. We can express the water loss due to the control error in terms of the layer thickness decrease. This makes the relation between the control tolerance, the layer thickness, and the process time explicit.

The ideal gas law relates the concentration *c* to the pressure and the temperature.

$$c = \frac{P}{RT}$$

Here the universal gas constant $R = 8.3\, JK^{-1}mol^{-1}$ and saturated vapor pressure $P = 2339Pa$ are used. We take the derivative of the concentration with respect to temperature.

$$\frac{dc}{dT} = -\frac{P}{RT^2} + \frac{1}{RT}\frac{dP}{dT}$$

This derivative of the pressure with respect to temperature is given by the Clausius-Clapeyron relation.

$$\frac{dP}{dT} = \frac{\Delta h}{T\Delta v}$$

Here, the change in specific latent heat upon evaporation $\Delta h = 2.5 \cdot 10^6\, J\, kg^{-1}$ and the change in specific volume upon evaporation $\Delta v = 57.8\, m^3 kg^{-1}$ are used to determine the evaporation rate.

Determination of the mass flux by the ideal gas law as described above results in an evaporation rate of $j = 4.4 \cdot 10^{-3}\, mol\, m^{-2}K^{-1}s^{-1}$. Multiplication with the molar mass of water and dividing this mass loss by the density will result in the rate of layer thickness decreases.

$$u_h = j \cdot M/\rho = 7 \cdot 10^{-8} m\, s^{-1}K^{-1}$$

Where $M = 18\, g\, mol^{-1}$ is the molar mass and $\rho = 1000\, kg\, m^{-3}$ is the density of water. This indicates that for every Kelvin of dewpoint error, the layer thickness decreases by 70 nm per second.

As shown above, the dewpoint temperature is 1.2 K below the chamber temperature when working at 20 °C and 93% humidity. When aiming for layer thicknesses in of only a few tens of nanometers, the concentrations and osmolarity of sample can change drastically if the sample is not cooled down to its dewpoint temperature.

Besides that, we can determine the temperature difference between the autogrid ring and the center of the grid. As the autogrid is cooled through the ring, the periphery of the grid will be at a lower temperature compared to the center of the grid. The ratio of thermal conductivity of gold over the thermal conductivity of air is about 1000, divided by the thickness ratio of about 150. For the thickness ratio, only half of the diameter was used because heat can diffuse in from both sides. The temperature difference between the air and the grid is 6 times larger than the temperature variation over the grid. This would mean that the temperature at the center of the grid is 0.2 degrees higher even if the edge is held perfectly at the dewpoint, which results in a decrease of layer thickness of 14 nm/s.
