## Supplementary figures and images for "Automated cryo-EM sample preparation by pin-printing and jet vitrification"

### Supplementary Figure 1

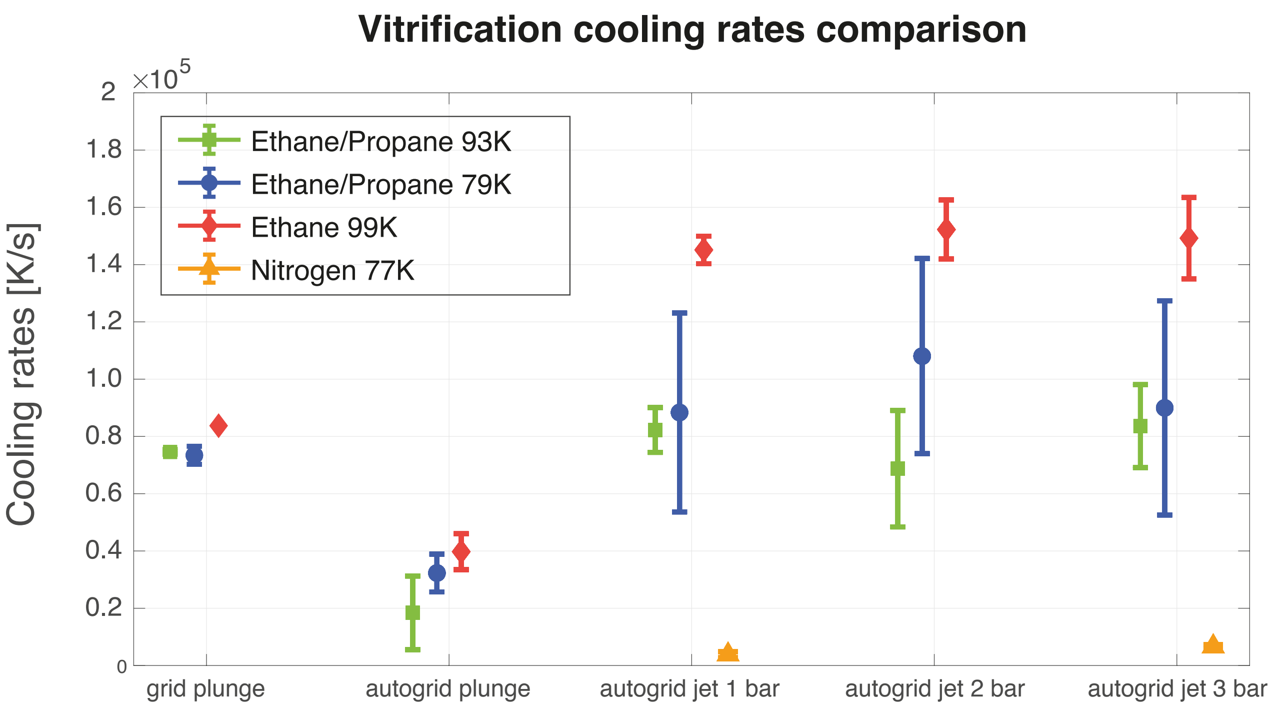


Supplementary Figure 1
